## Supplementary information for "Plectin-mediated cytoskeletal crosstalk as a target for inhibition of hepatocellular carcinoma growth and metastasis"

**1. Supplemental Material and Methods**
**2. Supplemental Figures & Supplemental Figure Legends**
**3. Supplemental Video Legends**
**4. Supplemental Table**
**5. Supplemental References**

### Supplemental Materials and Methods

#### Patient tissue samples

Formalin-fixed paraffin-embedded (FFPE) human liver tissue specimens were prepared at the Department of Surgery of the University Hospital Mannheim. The cohort consisted of 21 patients diagnosed with HCC (for details, see Supplementary file 1). Tissue collection and analysis were performed in accordance with institutional review board guidelines (reference no. 2012-293N-MA), and written informed consent was obtained from all included patients.

#### Animals

Liver-specific deletion of the plectin (Plec) gene was achieved by crossing *Plectin*$^{flox/flox}$ mice (*Ple*$^{fl/fl}$)[1] with *Alb-Cre* transgenic mice (MGI 2176228; The Jackson Laboratory, Bar Harbor, ME) to generate *Plectin*$^{lox/lox/Alb\text{-}Cre}$ (*Ple*$^{\Delta Alb}$) mice[2]. Immunodeficient NOD.Cg-Prkdcscid Il2rgtm1Wjl/SzJ (NSG) mice were purchased from the Czech Centre for Phenogenomics (BIOCEV – Institute of Molecular Genetics Academy of Sciences, Prague, Czechia)

Animals were housed under specific pathogen-free conditions with regular access to chow and drinking water and 12 h light/12 h dark conditions. All animal studies were performed in accordance with European Directive 86/609/EEC and were approved by the Czech Central Commission for Animal Welfare. Age-matched littermate mice were used in all experiments. The details regarding animal treatments can be found in the following sections.

#### Xenograft tumorigenesis

6-week-old NSG male mice were kept in a pathogen-free environment, and procedures were performed in a laminar airflow cabinet. $2.5\times10^6$ Huh7 cells in 250 µl PBS mice were administered subcutaneously in the left and right hind flanks. Animals were treated by orogastric gavage with plecstatin-1 every other day at a dose of 30 mg/kg. Before administration, plecstatin-1 was dissolved in olive oil. Tumor development was monitored by caliper measurement over 28 days before the mice were sacrificed and tissue samples were collected.

#### DEN-induced HCC mouse model

2-week-old *Ple*$^{fl/fl}$ and *Ple*$^{\Delta Alb}$ mice received intraperitoneal injection of 25 mg/kg diethylnitrosamine (DEN; Sigma-Aldrich, St. Louis, MO, USA) diluted in PBS. Mice were monitored for tumor formation 30 and 42 weeks after the DEN injection by magnetic resonance imaging (MRI), and tumor volumes were calculated from MRI images (for details see Magnetic resonance imaging section). Mice were sacrificed at 44 weeks post-injection, livers were dissected and tumors were measured using a caliper.

**Lung colonization assay**

Huh7 and SNU-475 cell lines stably expressing Red Firefly Luciferase reporter and GFP were prepared by lentiviral transfection of LentiGlo™ pLenti-CMV-RedFluc-IRES-EGFP plasmid (LP-31, Targeting Systems, El Cajon, CA, USA) according to manufacturer´s protocol. Next, $2\times10^6$ Huh7 or SNU-475 cells suspended in serum-free Dulbecco's modified Eagle medium (DMEM, Sigma-Aldrich) were injected into the tail vein of 5-week-old NSG mice. Mice were monitored for survival, and bioluminescence imaging was used to test for the presence of lung metastasis after 5 weeks. Prior to imaging, mice were anesthetized with isoflurane and injected intraperitoneally with D-luciferin potassium salt (Promega, Madison, WI, USA). Ten to fifteen min after injection, luciferase activity was measured using LagoX (Spectral Instruments Imaging, Tuscon, AZ, USA).

To prepare lung clearing samples, mice were injected with heparin (500 UI, Zentiva, Prague, Czechia) and after 15 min, mice were sacrificed by cervical dislocation and then perfused with 20 ml of PBS (pH 7.4) followed by 30 ml of 4% (w/v) paraformaldehyde (PFA) in PBS via the left ventricle of the heart. The excised organs were post-fixed in 4% (w/v) PFA at 4 °C for 24 h. The lungs were washed with PBS twice for 3 h and one time for 16 h to remove PFA just before clearing. The fixed lungs were immersed in 50% (v/v) clear, unobstructed brain imaging cocktails and computational analysis (CUBIC-1; 35w% water, 25 w% urea, 25w% N,N,N′,N′-tetrakis(2-hydroxypropyl)ethylenediamine and 15w% Triton X-100; 1:1 mixture of $H_2O$ and CUBIC-1) for more than 12 h and further immersed in CUBIC-1 with gentle shaking at 37 °C for 14 days. CUBIC-1 solution was refreshed every 2 days during the procedure. After decolorization and de-lipidation, the lungs were washed with PBS three times for 2 h. Lung samples after decolorization and de-lipidation were subjected to indirect immunostaining with anti-GFP antibody (1:100, A-11122, Invitrogen, Waltham, MA, USA) diluted in the staining buffer composed of 0.5% (v/v) Triton X-100, 0.4% BSA (BSA-1, Capricorn Scientific GmbH, Ebsdorfergrund, Germany) and 0.01% sodium azide for 5 days at room temperature (RT) with gentle shaking. The stained samples with primary antibody were washed with PBS three times for 3 h at RT and subsequently incubated for 5 days with secondary antibody anti-rabbit-AlexaFluor-594 (1:400, A11037, Life Technologies, Thermo-Fisher Scientific, Carslbad, CA, USA) at RT with gentle shaking. The stained samples with secondary antibody were washed with PBS three times for 3 h at RT.

Lung lobes were imaged using a Carl Zeiss Lightsheet Z1 microscope (Carl Zeiss AG, Oberkochen, Germany) with 5×/NA 0.15 objective with digital zoom 2. Lasers of 488 nm Solid-state laser 30 mW and 561 nm Solid-state laser 20 mW, were used for acquisition. The images were visualized and captured with Imaris software (version 10.1.0, Bitplane AG, Schlieren, Switzerland). The number of GFP positive foci was analyzed using a custom-made macro in ImageJ (NIH, USA) created by Martin Capek, Imaging Methods Core Facility (IMCF), Institute of Molecular Genetics (IMG).

**HDTVi-induced HCC mouse model**

For hydrodynamic tail vein injections, a mixture of plasmid containing 5 µg/ml of px330-expressing Tp53 sgRNA, 5 µg/ml of pT3-EF1a MYC DNA (92046, Addgene, Watertown, MA, USA), and 0.5 µg/ml pCMV HSB2 sleeping beauty transponase was prepared in sterile 0.9% sodium chloride (NaCl). 7-weeks-old $Ple^{fl/fl}$ and $Ple^{\Delta Alb}$ mice were pre-warmed for 15 min using two infrared lamps (IL 11, Beuer GmbH, Ulm, Germany), placed in a restrainer (TV-RED-150_STD, Braintree Scientific INC., Braintree, MA, USA) and injected into the lateral tail vein with a total volume corresponding 10% of body weight over 5–7 s. All animals were monitored daily, and animal experiments were performed in compliance with all relevant ethical regulations described in the animal permit. After mice were sacrificed, livers and lungs were visually inspected, excised, and photographed. Tumor samples were taken to obtain protein and the remaining liver tissue was incubated in 4% PFA for at least 24 h for FFPE tissue preparation.

**Proteomics on mouse tissue - sample preparation**

PBS-washed, snap-frozen livers were cut on dry ice and an aliquot of approximately 30 mg was suspended in 100 µl of lysis buffer (64 mM DTT, 8 M urea, 20% SDS, 1 M TEAB at pH 7.55). Each sample was homogenized using an ultra-sonicator. The samples were centrifuged (5000 g, 10 min, 4 °C), and the supernatant was transferred to a labeled Eppendorf tube and denatured at 95 °C for 5 min. Lysed tissues were stored at –20 °C.

**Magnetic resonance imaging**

Mice were anesthetized by inhalation of 1-2% isoflurane (Aerrane, Sigma-Aldrich) in atmospheric air. Animals were kept on a water tube heating pad to maintain constant body temperature. Breathing rate (~80 bpm) was monitored (ECG Trigger Unit, Rapid Biomedical, Rimpar, Germany). Imaging experiments were performed on a 4.7T horizontal bore MRI experimental system (Bruker Biospec 47/20, Ettlingen, Germany). A self-made wide-tunable 1H volume surface coil was used, which was optimized for whole-body mouse MRI (coil diameter = 35 mm). The livers were localized using axial T2*-weighted gradient echo sequence (repetition time (TR) 112 ms; echo time (TE) 3.7 ms; flip angle 30 °; averages = 4; field of view (FOV) 35×35 mm; plane resolution = 137x137 µm; slice thickness = 1.5 mm; scan time 1 min 54 s). Axial 1H T1-weighted anatomical images were acquired by gradient echo sequence (repetition time (TR) = 130 ms; echo time (TE) = 3.7 ms; flip angle = 70 °; averages =32; field of view (FOV) 35×35 mm; plane resolution = 137x137 µm; slice thickness 0.75 mm; scan time 17 min 44 s) to cover the whole liver. The pulse program and data processing were managed by Paravision 4.0 (Paravision, Bruker); then all data were post-processed and evaluated in ImageJ.

**Cell culture**

The human Huh7, Hep3B, PLC/PRF/5, SNU-398, SNU-423, and SNU-475 HCC cell lines were maintained in DMEM supplemented with 10% fetal bovine serum (FBS, Gibco, Thermo-Fisher Scientific) and 1% penicillin-streptomycin (Sigma-Aldrich) in 5% $CO_2$/air humidified atmosphere at 37 °C.

**CRISPR-mediated targeting of plectin**

Plectin KO cell lines were generated using the CRISPR/Cas9 system, by targeting exon 6 of *plectin* (Ensembl Plec-205), which encodes the essential actin-binding domain (ABD) and is shared among plectin isoforms. Single DNA sequence coding gRNA targeting sequence (AGTTGTCGCATCGCAGGCCC) was designed. To generate ΔIFBD mutant cell lines, an array of crRNA targeting sequences within exon 32 flanking the IFBD (5'-ACCCTAACACGGAGGAGAACCTC-3' and 5'-CGCTCCCGTTCCTCCTCGGTGGG-3') linked with short spacer sequence was designed (5'-AgatACCCTAACACGGAGGAGAACCTCAATTTCTACTCTTGTAGATCGCTCCCGTTCCTCCTCGGTGGG-3'). DNA sequences coding guide RNA targeting sequences and crRNA array were synthesized and subcloned into a modified version of a px330-Cas9-Venus[2] using *BbsI* restriction site, where Cas9 was fused to monomeric Venus as a selection marker. For WT cell generation, the px330-Cas9-Venus vector lacking the guide RNA targeting sequence was used. The vectors were transiently transfected into cells using Lipofectamine® LTX with Plus™ Reagent (Life Technologies) according to the manufacturer's instructions. Cells were incubated with the transfection complexes for 48 h, followed by fluorescence-activated cell sorting (FACS). Single-cell clones were FACS sorted into 96-well plates, and plectin knockout or mutation was confirmed by DNA sequencing and immunoblot analysis.

**Plecstatin treatment in vitro**

Plecstatin-1 (PST) was synthesized at the Faculty of Chemistry, University of Vienna, as previously described[3]. To determine $IC_{50}$ of PST, Huh7 and SNU-475 cells were seeded at a density of $5 \times 10^4$ cells/ml in a 96-well plate and incubated under standard conditions overnight. Next, the PST was diluted with culture medium to the desired concentration and cancer cells were treated for 96 h.

The cell viability was measured by absorbance using (3-(4, 5-dimethylthiazolyl-2)-2, 5-diphenyltetrazolium bromide) (MTT) viability assay (Sigma-Aldrich) according to the manufacturer´s protocol.

If not stated otherwise, PST was used at a concentration of 8 µM for 4 h. Stocks were dissolved in dimethylsulfoxide to a concentration of 4 mM and diluted in complete medium.

**Anchorage-independent growth in soft agar**

The wells of a 6-well plate were coated with 1.5 ml of 0.5% low-gelling temperature agarose (Sigma-Aldrich) solution dissolved in a standard culture medium. After solidification, $2\times10^3$ cells were suspended in 1 ml of 0.35% agarose in culture medium seeded on top of 0.5% agarose and overlaid with 200 µl culture medium. Cells were cultured for 28 days at 37 °C and 5% $CO_2$, and culture medium was exchanged once a week. For PST treatment, PST was added to 0.35% agarose mixture and into the overlaying medium in a final concentration of 8 µM. The colonies were stained and fixed with 0.01% crystal violet in 4% PFA in PBS and captured by Carl Zeiss Axio.Zoom V.16 microscope with PlanNeoFluar Z 1.0×/0.25 NA objective. The size and area of colonies were analyzed from binary masks created using a custom-made ImageJ macro.

**Micropatterning**

Cells were seeded on coverslips with crossbow-shaped micropatterns (CYTOOchips CW-L-A x18, CYTOO, Grenoble, France) according to the manufacturer´s protocol. In brief, micropatterns were coated with fibronectin overnight at 4 °C. After washing with PBS and complete medium, $3\times10^4$ cells were plated on coverslips, placed into 6-well plates, and allowed to adhere for 10 h before fixation. For PST treatment, PST was added 6 h after seeding to a final concentration of 8 µM. Cells were fixed with 4% formaldehyde (FA) in PBS for vimentin analysis or with 4% PFA in PBS for actin stress fiber analysis for 10 min at RT and permeabilized with 2% Triton in PBS for 5 min. Cells were imaged using a confocal Leica sp8 microscope (Leica Microsystems, Wetzlar, Germany) with HC PL APO 63×/1.4 NA immersion oil objective. Images were acquired as Z-stacks and deconvolved with Huygens Essential 4.0.0 software (Huygens; Scientific Volume Imaging, Hilversum, The Netherlands) using blind deconvolution. In vimentin networks analysis, the bundles were defined as a prominent signal of thickened vimentin filaments of more than approx. 0.5 µm, the clumps present condensed vimentin signal no longer resembling filamentous pattern. The vimentin fluorescence intensity distribution and focal adhesion distribution were analyzed using custom-made ImageJ macros. The peripheral region of the cell was defined as a 3 µm-wide border of the cell edge.

**Scratch wound-healing assay**

$1\times10^5$ Huh7 or SNU-475 cells were plated onto a well of 24-well plates in triplicate and incubated under standard conditions overnight to achieve 100% confluence. Prior to imaging, PST was added to a final concentration of 8 µM for 4 h. Using a 200-µl pipette tip, a linear scratch was made, running the length of each well. Subsequently, cells were washed with PBS and complete medium. To assess collective cell migration kinetics, 24-h movies for Huh7 and 18-h movies for SNU-475 with a 20-min time-lapse interval were recorded using brightfield video microscopy obtained with a Leica DMi8 with HCX PL FLUOTAR L 20×/0.40 NA objective (37 °C and 5% $CO_2$ in a humidified atmosphere; OKOlab, Naples, Italy). Individual fields of view were monitored in each well. The migration speed was calculated from the change of wound area between the initial and final timepoint, measured using ImageJ.

**Morphology in 3D collagen**

SNU-475 cells were trypsinized, washed in complete medium, and counted. $1\times10^5$ cells were mixed with 500 µl of 3 mg/ml Collagen R in complete medium. For PST treatment, the collagen mixture included PST at a final concentration of 8 µM. The suspension of cells in collagen (500 µl) was loaded onto a well of a 12-well plate, the gel was allowed to polymerize for 30 min at 37 °C and was subsequently overlaid with complete medium. After 24 h the cells were fixed with 4% PFA and stained with phalloidin, nuclei were counterstained with DAPI. The morphology of cells in 3D collagen was inspected using a Leica DMi8 microscope.

**Single cell random and chemotaxis migration assays**

SNU-475 cells were seeded at a total number of $1\times10^3$ onto a well of 6-well plate. Recordings of migration started 16 h after plating, and frames were taken with a Leica DMi8 with HCX PL FLUOTAR L 20×/0.4 NA objective in 5-min intervals over 20 h.

For chemotaxis experiments, $5\times10^3$ SNU-475 cells were seeded into Chemotaxis µ-slide (ibidi GmbH, Gräfelfing, Germany) according to the manufacturer´s protocol. Briefly, cells were incubated in 1% FBS-containing DMEM for 8 h after seeding, and chemo-attractant medium contained 20% FBS and 100 nM EGF. For plecstatin treatment, cells were treated with 8 uM PST 4 h after seeding.

The representative cell contours were generated by segmentation using ImageJ. The motion of cells was tracked using the ImageJ plugin Manual Tracking and the tracking results were analyzed using the ibidi Chemotaxis and Migration tool to generate Spider plots and processive indexes. The angle between cell protrusion and the direction of future cell motion was analyzed using a custom-made macro, using cell trajectories from Manual tracking analysis as an input for future movement calculation.

#### Traction force microscopy

Fibronectin-coated 7% polyacrylamide (PAA) gels (Young's modulus 25.5 kPa, thickness 130 µm), with embedded 0.2-µm fluorescent beads were prepared as described previously[4]. Briefly, the glass-bottom dishes were pre-treated with the 1:1:14 solution of acetic acid, bind-silane, and ethanol for 15 min at RT, washed with 95% ethanol, $H_2O$ and dried. Poly-L-lysine coated coverslips were incubated with 150 µm of 1:500 Fluorescent beads/mQ$H_2O$ (505/515, Invitrogen) for 20 min at 4 °C, subsequently washed with mQ$H_2O$ and dried. 11.8 µl of PAA gel was applied to the glass-bottomed dish, covered with a bead-coated coverglass, and incubated for 4 h at RT. After coverslip removal with tweezers and PBS wash, the gel was treated with 2 mg/ml Sulfo-SANPAH/$H_2O$ for 5 min under UV light and coated with 10 µg/ml fibronectin overnight at 4 °C.

$3\times10^4$ SNU-475 cells were seeded on the gels and incubated under standard conditions overnight. Prior to imaging, PST was added at a final concentration of 8 µM for 4 h. TFM imaging was performed using a Dragonfly (Andor Technology Ltd., Belfast, UK) spinning disc confocal microscope with Zyla 4.2 PLUS sCMOS camera and HC PL APO 63×/1.2 NA multi-immersion objective with water used as an immersion medium. During measurements, cells were kept in a life-cell imaging chamber (37 °C and 5% $CO_2$ in a humidified atmosphere; OKOlab). Gels were imaged both with the cells attached and in a relaxed state after cell trypsinization. Image acquisition was performed by Andor Technology software. Cell tractions were computed using pyTFM software[5] from images taken before and after trypsin-based cell removal.

#### Atomic force microscopy (AFM)

$2.5\times10^5$ Huh7 or SNU-475 cells were seeded onto fibronectin-coated glass coverslip placed in a 35 mm well 16 h before imaging. The cell elastic modulus was measured using AFM NanoWizard Sense (JPK Instruments AG, Berlin, Germany) with cantilever HYDRA-6R-200NG (AppNano, Mountain View, CA, USA; sensitivity = 76 nm/V, force constant = 30.13 N/m), with a spherical tip of 5.2 µm diameter. The cantilever was calibrated using the indentation of a microscope slide and using thermal noise. The measurement was performed using the force spectroscopy method in contact mode, using a sampling rate of 2046 Hz, a contact force of 4.186 nN, a loading speed of 1 µm/s, and a maximal indentation z-length of 5 µm. The cell elastic modulus was measured in an area of 16x16 µm, with a force map resolution of 8x8. The fitting of the force curves was performed using the Hertz-Sneddon model, and Young's modulus values were extracted using statistical parametric mapping (SPM) Data Processing software (JPK Instruments AG) when Poisson's ratio value of 0.5 was assumed. Young´s modulus of nuclear and cytosolic area was discerned based on the height profile, where the nucleus was defined as the area above 80% of the total cell height.

**Spheroid invasion assay**

$5\times10^5$ SNU-475 cells were grown for 2 days in Microtissues® 3D Petri Dish® (Sigma-Aldrich) according to the manufacturer's protocol. Formed spheroids were then embedded in 1.5 mg/ml Collagen I (Thermo-Fisher Scientific) solution containing DMEM, 5 % $NaHCO_3$ and 10% FBS. For PST treatment, PST was added to the collagen mixture and into the overlaying medium in a final concentration of 8 µM. Using a 96-well plate, one spheroid was embedded per well in 100 µl of collagen mixture, and once collagen polymerized, 80 µl of DMEM was added to the wells. Images were taken after 3 days using a Leica DMi8 microscope with HC PL FLUOTAR 5×/0.25 NA objective. The invaded area was calculated as the percentage of the initial spheroid area using ImageJ.

**Matrigel transwell invasion assay**

Matrigel Basement Membrane Matrix (Corning, Corning, NY, USA) diluted 1:3 in PBS was added to 6.5-mm transwell with 8-µm pore polycarbonate membrane inserts (Corning) and allowed to solidify at 37 °C for 2 h. Next, $3\times10^4$ SNU-475 cells in 180 µl total volume of serum-free medium were gently seeded on the top of the gel. For PST treatment, a serum-free medium mixture contained PST in a final concentration of 8 µM. The lower chamber was filled with 800 µl of medium containing 10% FBS as an attractant. After 20 h, cells were fixed using 4% PFA in PBS and stained with 0.02% crystal violet solution in 10% ethanol. Noninvasive cells in the upper matrix layer were discarded using a cotton swab and the cells on the bottom surface of the inserts were imaged using Leica DMi8 microscope with N PLAN 10×/0.25 NA objective.

**Gelatin degradation assay**

Dry coverslips were coated with a chilled poly-D-lysine (50 µg/ml) for 20 min at RT, washed 3 times with PBS, and subsequently coated with 0.5% glutaraldehyde for 15 min at RT. After washing 3 times with PBS, coverslips were coated with the mixture of 1% unlabeled gelatin and 488-conjugated gelatin (G13186, Thermo-Fisher Scientific) 1:30 for 30 min at 37 °C. $3\times10^4$ SNU-475 cells were seeded on fluorescent gelatin-coated coverslip placed onto a well of a 12-well plate in complete medium and allowed to adhere under standard conditions overnight. After 24 h, the coverslips were washed, fixed with 4% PFA in PBS for 10 min at RT, and permeabilized with 0.2% Triton X-100 in PBS for 5 min. F-actin was stained with Alexa Fluor 594-conjugated phalloidin (Thermo-Fisher Scientific), and nuclei were counterstained with DAPI. Cells were imaged using a Leica sp8 confocal microscope with HC PL APO 63×/1.40 NA immersion oil objective. 488-conjugated gelatin degraded area was segmented and measured using ImageJ.

**Matrigel invasion assay**

A biocompatible silicon self-stick cellular stopper (ibidi, 80209) was placed in the center of a 35-mm glass bottom dish (ibidi) before seeding $6\times10^5$ SNU-475 cells. After cells adhered, the stopper was removed and 250 µl of 50% BD Matrigel™ (4.5 mg/ml) in PBS was overlaid onto the cell monolayer to create a matrix barrier and allowed to polymerize for 2 h at 37 °C before adding growth medium on the top of the set Matrigel. Monolayers with overlaid Matrigel were then imaged with a Nikon time-lapse microscope or incubated for 24 h in a humidified atmosphere of 5% $CO_2$ at 37 °C before fixation and immunofluorescence. For time-lapse imaging, frames were taken using Leica DMi8 with HCX PL FLUOTAR L 20×/0.40 NA objective in 15-min intervals over 20 h. The motion of invading leader cells was tracked using the ImageJ plugin Manual Tracking. The angle between cell protrusion and the direction of future cell motion was analyzed using a custom-made ImageJ macro, using cell trajectories from Manual tracking analysis as an input for future movement calculation.

**Proteomic analyses of SNU-475 cell culture – sample preparation**

All steps for the sample collection process were performed on ice. $1.5\times10^6$ SNU-475 cells were seeded in P100 plates.

**Proteomics:** The cells were processed according to a nucleocytoplasmic fractionation protocol as previously described[3]. All steps were performed on ice. The medium was removed and the cells were washed twice with 1×PBS (3 mL). Then, PBS was thoroughly removed. Isotonic lysis buffer (10 mM HEPES, 10 mM NaCl, 3.5 mM MgCl2, 1 mM EGTA, 0.25 M Sucrose, 0.5% Triton X-100) containing protease inhibitors (1% PMSF, Sigma, and 1% protease and phosphatase inhibitor cocktail, Roche) was added to the wells, the cells were scraped off and transferred into labeled 15 mL Falcon tubes. The cellular membrane was ruptured using shear stress by pressing the cell suspension through a syringe 9–12 times. Membrane rupture with intact nuclei was monitored under a microscope. After centrifugation (960 g, 5 min), the supernatant containing the cytoplasmic (CYT) proteins was transferred into ice-cold ethanol (1:5, HPLC grade) and precipitated overnight at –20 °C. The pellet containing the nuclei was incubated with a hypertonic solution (10 mM Tris-HCl, 1 mM EDTA and 0.5 M NaCl) and subsequently with NP-40 buffer (10 mm Tris-HCl, 1 mm EDTA and 0.5% Triton X-100) containing protease inhibitors (1% PMSF, Sigma, and 1% protease and phosphatase inhibitor cocktail, Roche). Solubilization was assisted by an ultrasonic rod. After centrifugation (960 g, 5 min), the soluble nuclear (NE) proteins were also transferred into ice-cold ethanol (1:5, HPLC grade) and precipitated overnight at –20 °C. The precipitated proteins were finally pelleted by centrifugation (5000 g, 30 min, 4 °C), the solution was decanted and the pellet dried under vacuum.

**Phosphoproteomics:** The cells were washed twice with 2 ml cold Tris-buffered saline (TBS; 25 mM Tris·HCl, 150 mM NaCl, pH 7.4). The wash buffer was completely removed. A volume of 250 µl of 4% [wt/vol] sodium deoxycholate buffer (SDC; 0.4 g SDC, 500 µl Tris·HCl (pH 8.8) in 9.5 ml H2O) was added to the well. The cells were scraped off and the whole cell lysate (WCL) was transferred to an Eppendorf® tube, which was subsequently heated to 95 °C for 5 min. WCLs were stored at –20 °C until processing.

**Image acquisition and processing**

If not stated otherwise, immunofluorescence images were acquired at RT using a Leica DM6000 wide-field microscope (Leica Microsystems) with Leica DFC 9000 sCMOS camera and an APO 100×/1.4 NA immersion oil objective with Type F Immersion liquid (Leica Microsystems). The acquisition was performed by LasX software (Leica Microsystems). Immunofluorescence confocal images were acquired at RT using a Leica TCS SP8 confocal fluorescence microscope (Leica Microsystems) with photomultiplier tubes (PMTs), supersensitive hybrid detectors (HyDs), and HC PL APO 63×/1.4 NA immersion oil objective, using Type F Immersion liquid (Leica Microsystems). The acquisition was performed by LasX software (Leica Microsystems). Unless stated otherwise, representative maximal projections of z-stacks, acquired from whole cell volumes are shown. Post-acquisition processing (such as formation of maximal projections, linear adjustments of brightness and contrast, inversion of grayscale images, changes in lookup tables, cropping or resizing) was performed using an open-source ImageJ image processing package[6]. Soft agar colony number and size, focal adhesion distribution, vimentin fluorescence intensity distribution, and direction of protrusion formation, were analyzed using custom-made ImageJ macros created by Jan Valecka, IMCF IMG.

**Immunoblot analysis**

$7\times10^5$ cells were seeded on a P100 plate coated with fibronectin (Sigma-Aldrich) and incubated for 24 h. For PST treatment, PST in a final concentration of 8 µM was added 4 h after cell seeding. Cells were rinsed with PBS and lysed in RIPA buffer (20 mM Tris pH 7.4, 150 mM NaCl, 0.1% sodium dodecyl sulfate, 0.5% Na-Deoxycholate, 1% Triton X-100), supplemented with protease and phosphatase inhibitors (Roche, Basel, Switzerland). For liver tissue lysate preparation, snap-frozen liver tissue samples were homogenized in ice-cold RIPA buffer supplemented with a protease and phosphatase inhibitor cocktail (Roche) using the Tissue Lyzer II (Qiagen, Hilden, Germany). Protein concentrations were determined using a BCA Protein Assay Kit (Thermo-Fisher Scientific). Proteins were separated by SDS-PAGE and transferred to nitrocellulose membranes. Membranes were incubated with 5% BSA in PBST to block non-specific antigen interactions and subsequently incubated with primary and secondary antibodies (see Supplementary file 2). Signals were detected and quantified using an Odyssey imaging system (LI-COR Biosciences, Lincoln, NE; Figure 2—figure supplement 1C, Fig. 3D, Figure 3—figure supplement 1C; Figure 5—figure supplement 1C) or using chemiluminescence *via* enhanced chemiluminescence (ECL) western blot substrates (Thermo-Fisher Scientific; Fig. 1D,H).

**Immunohistochemistry and immunofluorescence of patient samples**

Immunohistochemistry and immunofluorescence microscopy of patient samples were performed by Danijela Heide and Jenny Hetzer at the Department of Chronic Inflammation and Cancer, DKFZ, (Heidelberg). 2-µm sections were cut from paraffin blocks with a Microtome (Leica Biosystems, Nussloch, Germany), mounted on glass coverslips (Thermo-Fisher Scientific), and H&E stained using a Leica BOND-MAX automated staining machine. Plectin immunofluorescence staining was done with primary antibody (Progen, Heidelberg, Germany; GP21, dilution 1:500) followed by the AKOYA Biosciences Opal Fluorophore kit (Opal 620, FP1495001KT diluted 1:150) and counterstained with AKOYA 10× Spectral DAPI (FP1490) following the manufacturer's instructions. The slides were imaged with a NanoZoomer S60 Scanner (Hamamatsu, Japan) at 20× and analyzed with the NDP.view software (ver. 2.7.25, Hamamatsu).

**Immunohistochemistry on murine samples**

FA-fixed liver paraffin sections (4 µm) sections were subjected to heat-induced antigen retrieval in Tris-EDTA (pH 9) or citrate (pH 6) buffer and further permeabilized with 0.1 M glycine, 0.1% Triton X-100 in PBS for 15 min. Blocking of unspecific antigen interactions was performed with 5% BSA in PBS. Next, sections were incubated with primary antibodies at 4 °C overnight followed by incubation with secondary antibodies at RT for 120 min, nuclei were counterstained with DAPI. To evaluate the apoptosis rate in liver tumor samples, apoptotic cells were visualized on paraffin-embedded liver tissue sections using a Click-iT TUNEL Alexa Fluor 488 kit (ThermoFisher Scientific) according to the manufacturer's protocol. Immunofluorescence images were acquired using DM6000 (Leica Microsystems; with HC PL Apo 10×/0.40 NA and PH1 HC PLAN Apo 20×/0.70 NA objectives) and analyzed using ImageJ.

**Patient analysis – data mining**

To investigate the association of *plectin* mRNA expression with gene signatures and clinical parameters in HCC patients, mRNA expression datasets were obtained from online repositories including Gene Expression Omnibus (GEO), ArrayExpress, and the Cancer Genome Atlas. For Affymetrix whole-genome microarray data, raw .CEL files were processed using the SCAN (single channel array normalization) method[7], and samples with median GNUSE quality scores[8] above 1.25, as computed by the "frma" Bioconductor package, were excluded. For all other datasets, processed data from the original authors were used. Cases of fibrolamellar carcinoma were removed[9]. Across all platforms, probe or gene IDs were mapped to Entrez gene identifiers. For microarray platforms containing multiple alternative probes or probesets per gene, the probe showing the great variance in expression among HCC samples was selected. Meta-analyses across datasets were computed with the "metafor" R package[10] using a random-effects model and the DerSimonian-Laird estimator. The tumor node metastasis (TNM) analysis comprised datasets corresponding to Iizuka et al.[11], GSE14520, iCOD[12], GSE76427, GSE36376, and TCGA-LIHC from Affymetrix HuGeneFL, Affymetrix HT_HG-U133A, and Affymetrix HG-U133 Plus 2.0, Illumina HumanHT-12 V4.0, and Illumina HiSeq platforms. Collections of gene signatures were obtained from the Molecular Signatures Database (MSigDB)[13,14], and used to produce quantitative gene set enrichment scores with the "GSVA" (Gene Set Variation Analysis) Bioconductor package[15]. To assess the correlation of *PLEC* expression with various genes and pathways in human HCC, eleven datasets with extensive whole-genome RNA expression data were chosen (TCGA-LIHC[16], GSE65485, GSE50579, GSE45436, GSE62232, GSE9843[17], iCOD[18], GSE63898, GSE64041[19], GSE76297, GSE16757), and gene expression values were batch-adjusted using the ComBat method[20] as implemented by the "sva" R package. To generate t-SNE plots, the "Rtsne" package was applied to ComBat-adjusted data with a perplexity parameter of 50. The Nearest Template Prediction method[21] was used to assign HCC samples to the most likely predicted molecular subclass based on published gene lists[16,17,19,22,23]. The Kaplan-Meier survival analysis of *PLEC* expression was based on three datasets for which recurrence-free survival data was available: GSE17856, GSE14520, and GSE76427.

**Proteomics on mouse tissue samples**

Samples were thawed and the protein concentration was determined via bicinchoninic acid (BCA) assay and an aliquot of 20 µg of protein was used for proteomics. Proteins were reduced with dithiothreitol (DTT, 64 mM) and carbamidomethylated with iodoacetamide (148 mM). Proteolytic digestion was performed using the ProtiFi S-trap technology and samples were loaded onto S-trap mini cartridges. Trapping buffer (methanol 90% v/v, 0.1 M tetraethylammonium bromide was added.

Samples were thoroughly washed and digested with 0.5 µg trypsin/Lys-C mix (Promega) at 37 °C for 2 h (1:40 enzyme-to-protein ratio). Peptides were eluted, dried in glass inserts, and stored at -20 °C. Dried samples were reconstituted with 5 µl of a peptide standard mix consisting of the four synthetic peptides at 10 fmol in 30% formic acid (Glu1-fribrinopeptideB, EGVNDNEEGFFSAR; M28, TTPAVLDSDGSYFLYSK; HK0, VLETKSLYVR and HK1, VLETK(ε-Ac)SLYVR) and diluted with loading buffer (40 µl, acetonitrile 2% v/v, TFA 0.05% v/v; ddH2O 97.95% v/v). The glass inserts were transferred into 2 ml Eppendorf tubes filled with 400 µl ddH2O. The samples were sonicated for 5 min (20% power), centrifuged (5000 g, 7 min, RT), and transferred into labeled high-performance liquid chromatography vials.

LC-MS/MS data acquisition was performed employing a timsTOF Pro mass spectrometer (Bruker Daltonics, Bremen, Germany) hyphenated to a Dionex UltiMate™ 3000 RSLCnano system (Thermo Scientific, Bremen, Germany). Samples were analyzed in data-dependent acquisition mode by label-free quantification (LFQ) shotgun proteomics in PASEF mode[24]. Samples were loaded on an Acclaim™PepMap™ C18 HPLC pre-column (2 cm × 100 µm, 100 Å, Thermo-Fisher Scientific) at a flow rate of 10 µl/min. After trapping, peptides were eluted at a flow rate of 300 nL $min^{-1}$ and separated on an Aurora series CSI UHPLC emitter column (25 cm × 75 µm, 1.6 µm C18, Ionopticks, Fitzroy, Australia) applying an 8–40% gradient of mobile phase B (79.9% ACN, 20% $H_2O$, 0.1% FA) in mobile phase A (99.9% $H_2O$, 0.1% FA). The injection volume was 2 µl. The gradient for the proteome profiling was selected to run over 85 min giving a total run time of 135 min per sample.

Protein identification was performed *via* MaxQuant (version 1.6.17.0) employing the Andromeda search engine against the UniProt Database (mus musculus, version 06/2021, 17'519 entries). A mass tolerance of 20 ppm for MS spectra and 40 ppm for MS/MS spectra, a PSM–, protein– and site-false discovery rate (FDR) of 0.01, and a maximum of two missed cleavages per peptide were allowed. Match-between-runs was enabled with a matching time window of 0.7 min and an alignment time window of 20 min. Oxidation of methionine and N-terminal protein acetylation were set as variable modifications. Carbamidomethylation of cysteine was set as a fixed modification. Data analysis was performed *via* Perseus software (version 1.6.14.0)[25]. Proteins with at least a 75% quantification rate in at least one group were considered for analysis. A two-sided Student's *t*-test with S0 = 0.1 and FDR cut-off of 0.05 was applied for identifying multiple testing-corrected significantly regulated proteins.

**Proteomic analyses of SNU-475 cell culture**

**Proteomics:** The protein fractions were pelleted, dried and solubilised in sample buffer (7.5 M urea, 1.5 M thiourea, 4% CHAPS, 0.05% SDS and 100 mM dithiothreitol) before the total protein amount was determined by means of a colorimetric Bradford assay. A total of 20 µg protein per sample were then tryptically digested. This involved reduction with dithiothreitol, alkylation with iodoacetamide and digestion with trypsin/Lys-C according to a filter-assisted protein digestion (FASP) protocol[3]. The obtained peptide samples were dried in a speed-vac.

**Phosphoproteomics:** WCLs were thawed and lysed using the S220 Focused-ultrasonicator (Covaris, LLC., Woburn, MA, USA). Protein concentrations were determined via bicinchoninic acid assay (BCA)-assay. Between 130–220 µg of protein was reduced and alkylated with tris(2-carboxyethyl)phosphine (TCEP) and 2-chloroacetamide (2-CAM) for 5 min at 45 °C, followed by 18 h digestion with Trypsin/Lys-C (1:100 enzyme-to-substrate ratio) at 37 °C. An aliquot of 20 µg peptide was taken from each sample for global proteome analysis and dried in a vacuum concentrator. Then, the samples were reconstituted in styrenedivinylbenzene-reverse phase sulfonate (SDB-RPS) loading buffer (99% iPrOH, 1% TFA) and desalted via SDB-RPS StageTips. Desalted global proteome samples were reconstituted in 5 µL formic acid (30%) containing synthetic standard peptides at 10 fmol and diluted with 40 µL loading solvent (98% H2O, 2% ACN, 0.05% TFA). For phosphopeptide enrichment, the digested samples were mixed with enrichment buffer (52% H2O, 48% TFA, 8 mM KH2PO4) and incubated with TiO2 Titansphere beads (GL Sciences, Japan), followed by sample clean-up via C8-StageTips. Phosphopeptides were eluted and subsequently dried in a vacuum concentrator. Phosphopeptides were reconstituted in 15 µL MS loading buffer (97.7% H2O, 2% ACN, 0.3% TFA).

LC-MS/MS analyses were performed employing a timsTOF Pro mass spectrometer (Bruker Daltonics, Bremen, Germany) hyphenated with a Dionex UltiMateTM 3000 RSLCnano system (Thermo Scientific, Bremen, Germany). Samples were analyzed in data-dependent acquisition mode by label free quantification (LFQ) shotgun proteomics. The injection volume was 10 µL and 2 µL for analyzing phosphoproteomes and global proteomes, respectively. Samples were loaded on an AcclaimTMPepMapTM C18 HPLC pre-column (2 cm × 100 µm, 100 Å, Thermo Fisher Scientific™, Vienna, Austria) at a flow rate of 10 µL min-1 MS loading buffer. After trapping, peptides were eluted at a flow rate of 300 nL min-1 and separated on an Aurora series CSI UHPLC emitter column (25 cm × 75 µm, 1.6 µm C18, Ionopticks, Fitzroy, Australia) applying a gradient of 8–40% mobile phase B (79.9% ACN, 20% H2O, 0.1% FA) in mobile phase A (99.9% H2O, 0.1% FA) over 95 min.

Protein identification was performed via MaxQuant (version 1.6.17.0) employing the Andromeda search engine against the UniProt Database (version 11/2021, 20'375 entries). A mass tolerance of 20 ppm for MS spectra and 40 ppm for MS/MS spectra, a PSM–, protein– and site-false discovery rate (FDR) of 0.01, and a maximum of two missed cleavages per peptide were allowed. Match-between-runs was enabled with a matching time window of 0.7 min and an alignment time window of 20 min. Oxidation of methionine, N-terminal protein acetylation, and phosphorylation of serine, threonine, and tyrosine were set as variable modifications. Carbamidomethylation of cysteine was set as a fixed modification. Global proteome data analysis was performed via Perseus (version 1.6.14.0). Proteins with at least 60% quantification rate in at least one group were considered for analysis. A two-sided Student's t-test with S0 = 0.1 and an FDR cut-off of 0.05 was applied for identifying multiple testing-corrected significantly regulated proteins or phosphosites.

**Statistical analyses**

Proteomics on mouse tissue samples proteomics of SNU-475 cell cultures (see details in corresponding sections), all graphs and statistical tests were performed using GraphPad Prism (GraphPad Software, Inc., La Jolla, CA). Rose graphs were generated using Microsoft Excel (Microsoft Corporation, Redmont, WA). In the boxplots the box margins represent the 25th and 75th percentile with the median indicated; whiskers reach the last data point. Data comparison of adjacent tumor and non-tumor tissue was performed using a paired *t*-test. Data comparison of individual experimental groups with the control group was performed using a two-tailed *t*-test. Growth curves were analyzed using two-way ANOVA. Survival curves were analyzed using the Mantel-Cox test. Data distribution was assumed to be normal, but this was not formally tested. Statistical significance was determined at the level of $^*P < 0.05$, $^{**}P < 0.01$, $^{\dagger}P < 0.001$. The number of independent experiments (N), number of data points (n), and statistical tests used are specified for individual experiments in the figure legends.

**Antibodies and reagents**

For details on primary and secondary antibodies used in this study, see Supplementary file 2. The following reagents were used: AF-Phalloidin 594 (Invitrogen); low gelling temperature agarose (Sigma-Aldrich), Collagen I Rat Protein (Thermo-Fisher Scientific), Crystal Violet solution (Sigma-Aldrich), DAPI (AppliChem, Darmstadt, Germany), diethylnitrosamine (DEN; Sigma-Aldrich), mouse EGF recombinant protein (Invitrogen), Gelatin 488 Conjugate (Thermo-Fisher Scientific), Fibronectin (FN) from bovine plasma (Sigma), glutaraldehyde solution 25% (Sigma-Aldrich), heparin (Zentiva), isoflurane (Aerrane, Sigma-Aldrich), D-luciferin potassium salt (Promega), Matrigel Basement Membrane Matrix (Corning), Poly-D-Lysine solution (Sigma-Aldrich), Sulfo-SANPAH (Sigma-Aldrich), N,N,N′,N′-Tetrakis (2-Hydroxypropyl)ethylenediamine (Sigma-Aldrich).

### Supplemental Figures & Supplemental Figure legends

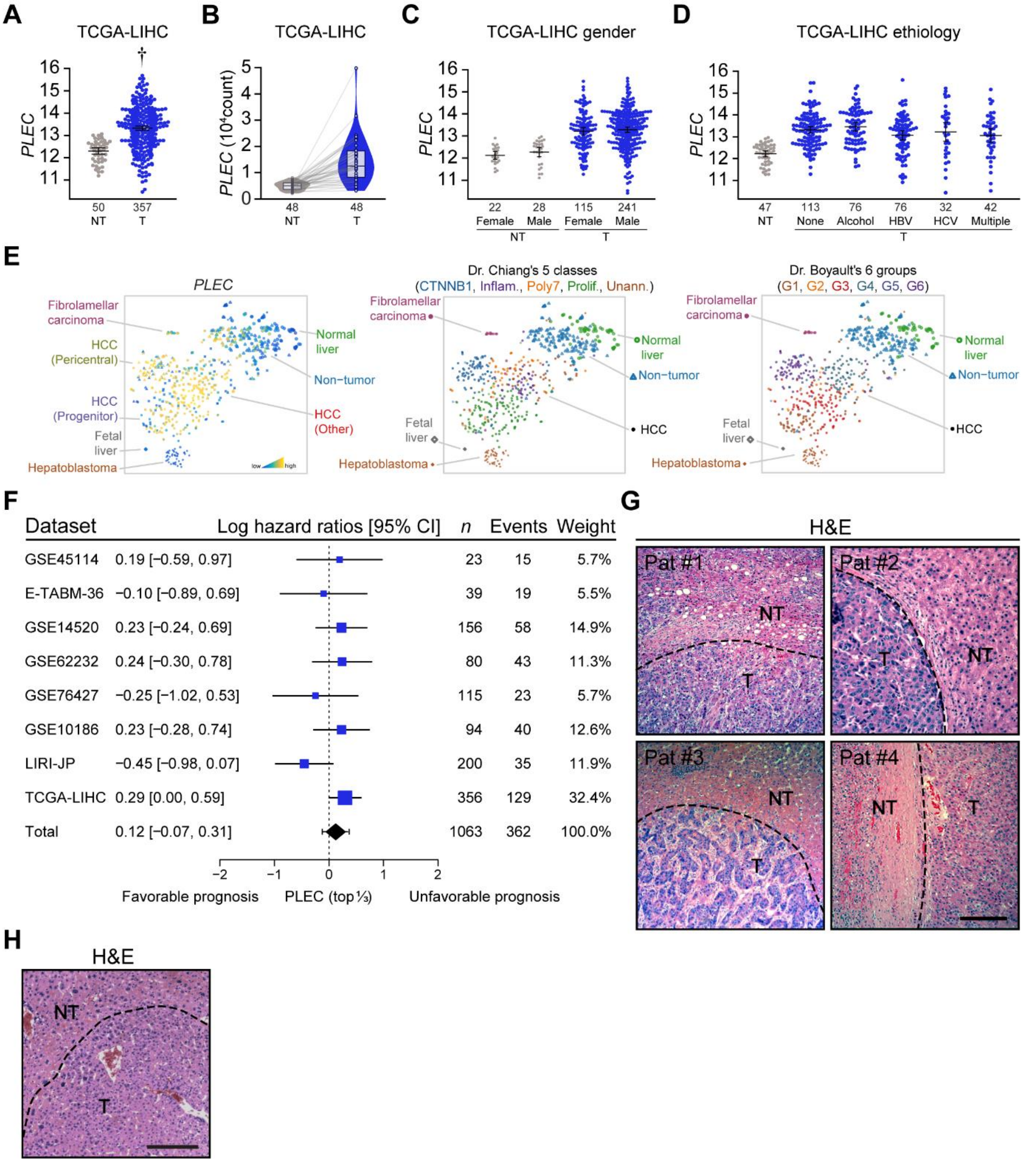


**Figure 1—figure supplement 1. Plectin is elevatated in HCC across genders and etiologies. (A-D)** TCGA-based analysis of *plectin* (*PLEC*) mRNA expression in non-tumor (NT) and tumor (T) tissue of liver hepatocellular carcinoma (LIHC). Graphs show differential *plectin* expression in the whole cohort (A), pair-matched samples (B), and samples sorted by gender (C) and etiology (D). The numbers of included participants per cohort are indicated in the graph. **(E)** t-SNE plots (left graph) show *plectin* (*PLEC*) mRNA expression in subgroups of HCC patients. Points, individual patient tissue samples. As a reference, t-SNE plots of Dr. Chiang's (middle graph) and Dr. Boyault's (right graph) classification are

shown. The HCC classes were predicted using the Nearest Template Prediction method. **(F)** Meta-analysis of the predictive value of *plectin* expression across different HCC datasets. Blue squares indicate the log hazard ratios and 95% confidence interval from a Cox proportional hazards model. The black diamond represents the mean and 95% confidence interval for the overall log hazard ratio, while the whiskers indicate the 95% prediction interval. **(G)** Representative images of H&E-stained human HCC tissue sections corresponding to immunofluorescence images shown in Fig. 1C. NT, non-tumor area; T, tumor area. Scale bar, 200 µm. **(H)** Representative image of H&E-stained section of DEN-induced HCC corresponding to immunofluorescence images shown in Fig. 1G. NT, non-tumor area; T, tumor area. Scale bar, 200 µm.

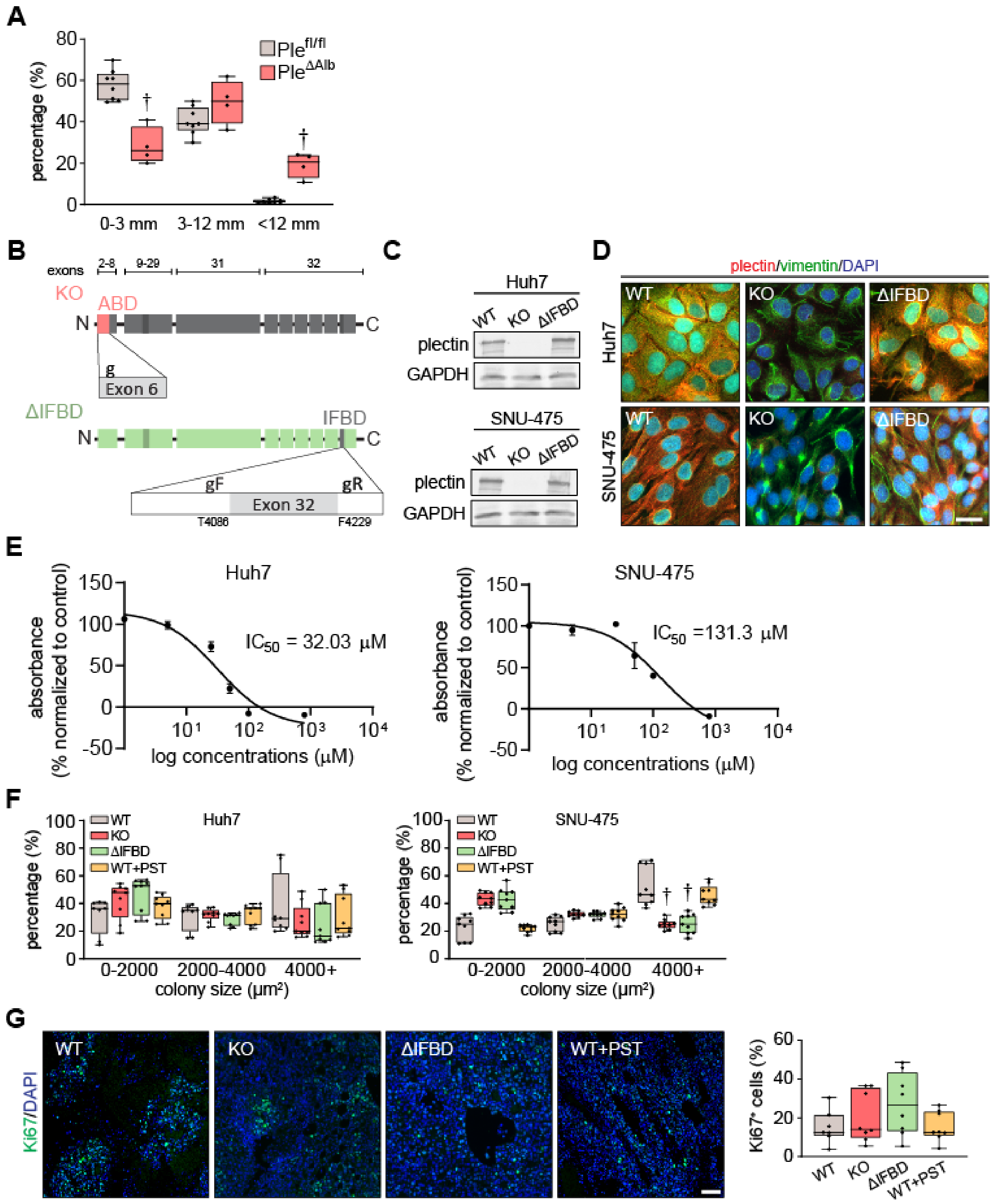


**Figure 2—figure supplement 1. Plectin influences HCC progression *in vivo*. (A)** Percentage of DEN-induced *Ple*$^{fl/fl}$ and *Ple*$^{\Delta Alb}$ HCCs corresponding to the indicated size intervals. Boxplot shows the median, 25th, and 75th percentile with whiskers reaching the last data point; dots, individual mice; $N = 8$ (*Ple*$^{fl/fl}$), 4 (*Ple*$^{\Delta Alb}$). Two-tailed *t*-test; †$P < 0.001$. **(B)** Schematic of CRISPR/Cas9-based strategy for the generation of plectin KO and ΔIFBD mutant HCC lines. To generate plectin KO, exon 6 of plectin was targeted by the single guide RNA (g) as previously described[26]. To generate the plectin ΔIFBD HCC lines, deletion between amino acid T4086-F4229 (Uniprot accession: Q15149-9) was introduced using

crRNA array targeting the sequences within the exon 32 flanking the IFBD. Upper bar indicates the exons encoding plectin domains shown below. For details, see the Materials and methods section. **(C)** Representative immunoblots for plectin in WT, KO, and ΔIFBD Huh7 and SNU-475 cell lines. GAPDH, loading control. **(D)** Representative images of WT, KO, and ΔIFBD Huh7 and SNU-475 cells immunolabeled for plectin (red) and vimentin (green). Nuclei, DAPI (blue). Scale bar, 10 µm. **(E)** $IC_{50}$ curve of Huh7 and SNU-475 cells after treatment with plecstatin-1 (PST) for 96 hours. **(F)** Percentage of colonies grown from WT, KO, ΔIFBD, and PST-treated WT (WT+PST) Huh7 and SNU-475 cells within indicated size intervals. Boxplots show the median, 25th, and 75th percentile with whiskers reaching the last data point; dots, agar wells; $n$ = 9 agar wells; $N$ = 3. Two-way ANOVA; †$P$ < 0.001. **(G)** Representative images of WT, KO, ΔIFBD, and WT+PST Huh7 xenograft sections immunolabeled for Ki67 (green). Nuclei, DAPI (blue). Scale bar, 150 µm. Quantification (percentage) of Ki67-positive cells in xenografts grown from WT, KO, ΔIFBD, and WT+PST Huh7 cells shown in Fig. 2H. Boxplot shows the median, 25th and 75th percentile with whiskers reaching the last data point; dots, individual tumors; $N$ = 7 (WT), 8 (KO), 8 (ΔIFBD), 8 (WT+PST).

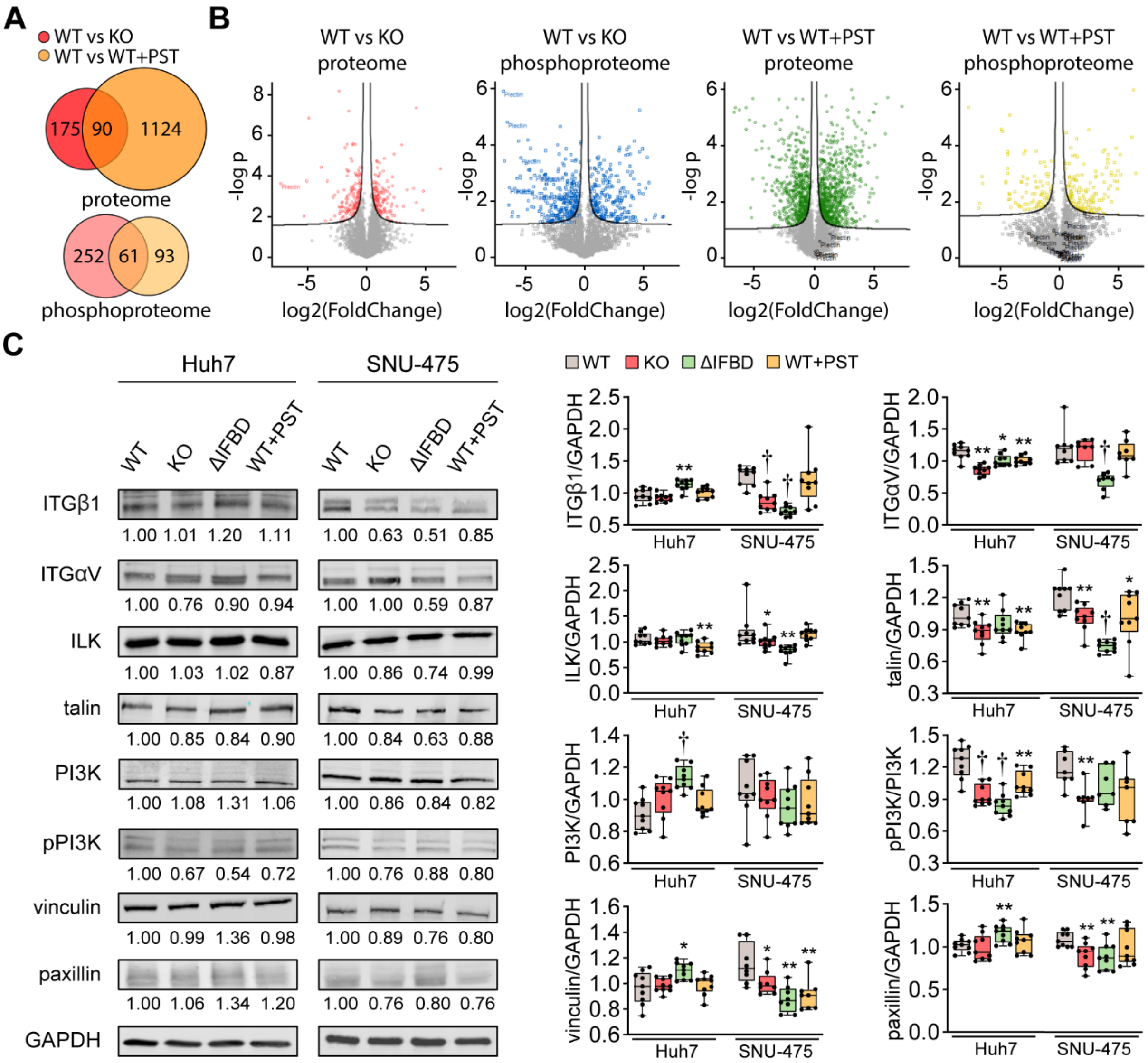


**Figure 3—figure supplement 1. Integrin-associated signaling is altered in SNU-475 cells upon CRISPR/Cas9- or PST-mediated plectin inactivation. (A)** Venn diagrams show relative proportions of differentially expressed/phosphorylated proteins identified by proteomic (proteome) and phospho-proteomic (phosphoproteome) analyses of WT vs KO and WT vs WT+PST SNU-475 cells shown in Fig. 3A-C. **(B)** Volcano plots show the fold change vs. *p*-value of differentially expressed/phosphorylated proteins of indicated comparisons of Snu475 cells. **(C)** Quantification of β1 integrin (ITGβ1), αV integrin (ITGαV), ILK, talin, PI3K, phospho-p85 (Tyr458)/p55(Tyr199)-PI3K (pPI3K), vinculin, and paxillin in indicated Huh7 and SNU-475 cell lines by immunoblotting. GAPDH, loading control. The numbers below lines indicate relative band intensities normalized to average WT values. Boxplots show relative band intensities normalized to GAPDH or non-phosphorylated protein. The box represents the median, 25th, and 75th percentile with whiskers reaching the last data point; dots, individual experiments; $N = 9$. Two-tailed *t*-test; *$P < 0.05$; **$P < 0.01$; †$P < 0.001$.

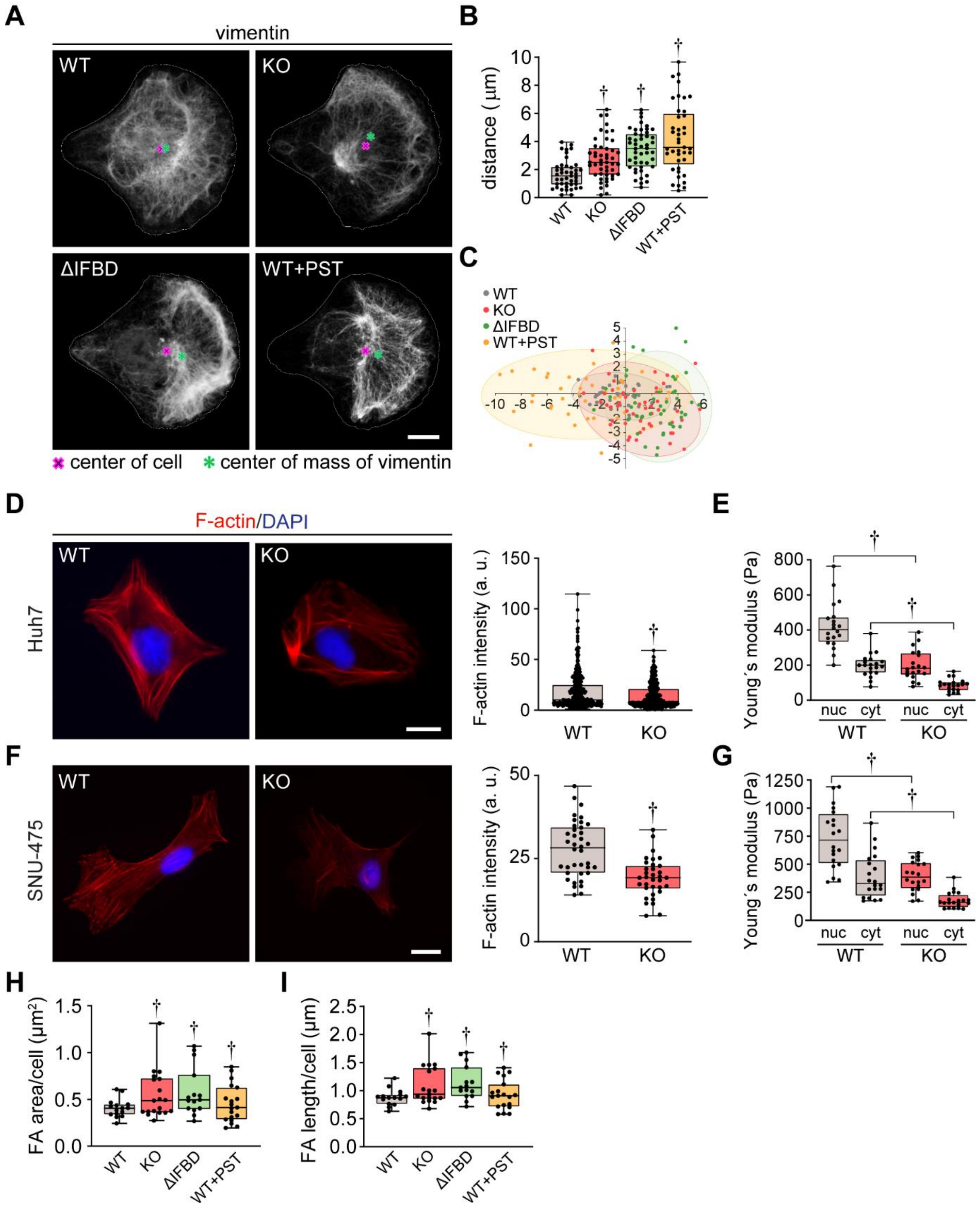


**Figure 4—figure supplement 1. Cytoskeletal networks and cell stiffness are altered upon plectin inactivation. (A)** Representative confocal images of micropattern-seeded WT, KO, ΔIFBD, and PST-treated WT (WT+PST) SNU-475 cells immunolabeled for vimentin. Line, cell contour; red cross, center of cell; green asterisk, center of mass of vimentin fluorescence signal. Scale bar, 10 µm. **(B,C)** Quantification of vimentin distribution in WT, KO, ΔIFBD, and PST-treated WT (WT+PST) SNU-475 cells shown in (A). Graphs show the distance between the cell center and the center of mass of the vimentin

signal (B) and the position of the center of mass of the vimentin signal (C). Boxplot shows the median, 25th and 75th percentile with whiskers reaching the last data point; dots, individual cells; $n$ = 43 (WT), 48 (KO), 47 (ΔIFBD), 41 (WT+PST) cells; $N$ = 3. Two-tailed $t$-test; †$P < 0.001$. **(D,F)** Representative images of WT and KO Huh7 (D) and SNU-475 (F) cells immunolabeled for F-actin (red). Nuclei, DAPI (blue). Scale bar, 20 µm. Graphs show the quantification of F-actin fluorescence intensity. Boxplots show the median, 25th, and 75th percentile with whiskers reaching the last data point; dots, individual cells; $n$ (Huh7) = 169 (WT), 180 (KO); $n$ (SNU-475) = 38 (WT), 34 (KO) cells; $N$ (Huh7) = 3; $N$ (SNU-475) = 3. Two-tailed $t$-test; †$P < 0.001$. **(E,G)** Quantification of average Young´s modulus values in nuclear (nuc) and cytosolic (cyt) area of WT and KO Huh7 (E) and SNU-475 (G) cells. Boxplot shows the median, 25th, and 75th percentile with whiskers reaching the last data point; dots, individual cells; $n$ = 20 cells; $N$ = 3. Two-tailed $t$-test; †$P < 0.001$. **(H,I)** Quantification of average FA length (H) and FA area (I) per cell for WT, KO, ΔIFBD, and PST-treated WT (WT+PST) SNU-475 cells shown in Fig. 4F. Boxplot shows the median, 25th, and 75th percentile with whiskers reaching the last data point; dots, individual cells; $n$ = 15 (WT), 18 (KO), 20 (ΔIFBD), 19 (WT+PST) cells; $N$ = 3. Two-tailed $t$-test; †$P < 0.001$.

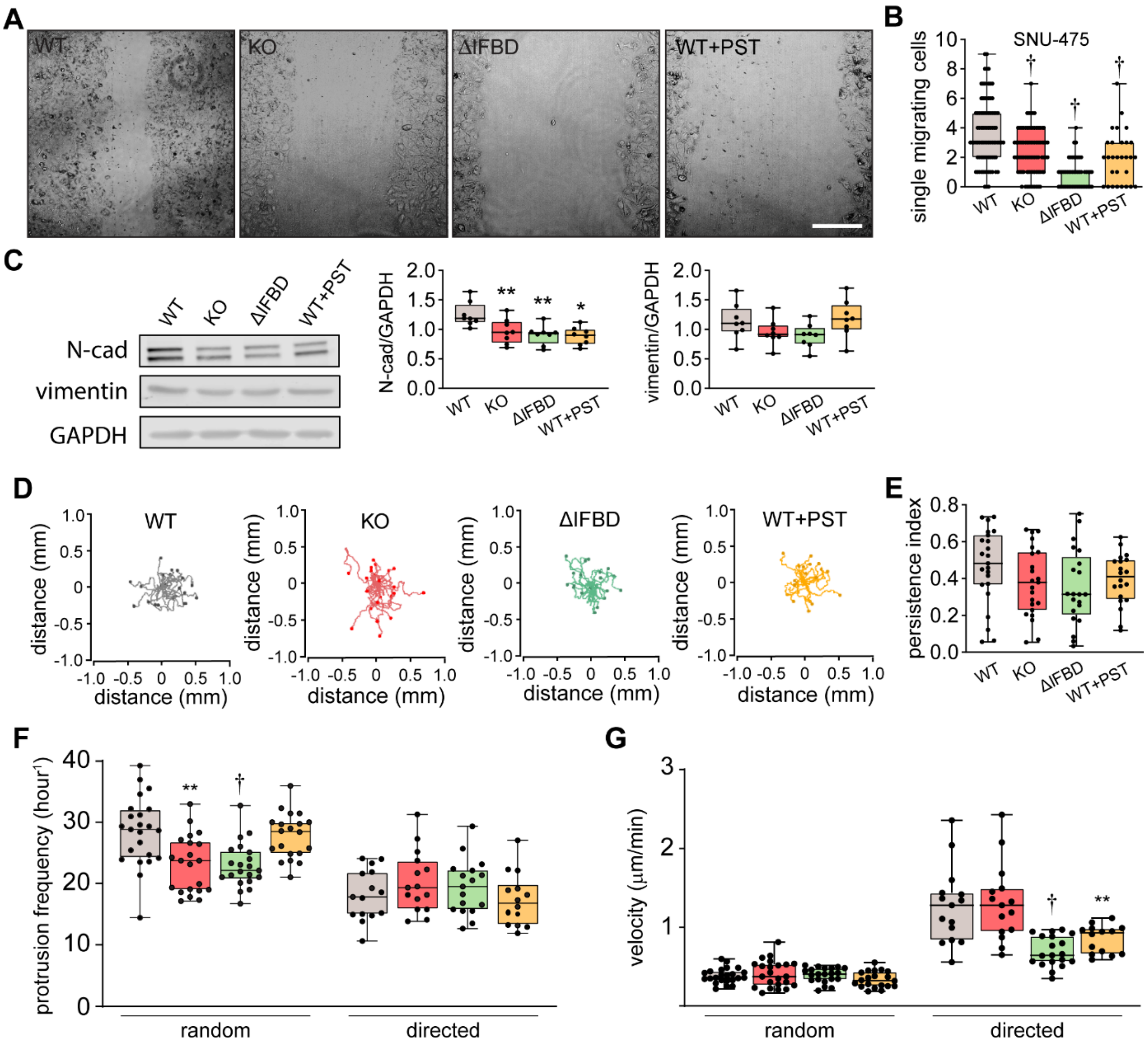


**Figure 5—figure supplement 1. Plectin inactivation impairs migration of HCC cells. (A)** Representative phase contrast images of WT, KO, ΔIFBD, and PST-treated WT (WT+PST) Huh7 cells migrating in the scratch-wound assay for 24 hours corresponding to the graph shown in Fig. 5B. Scale bar, 200 µm. **(B)** Quantification of the number of WT, KO, ΔIFBD, and PST-treated WT (WT+PST) leader SNU-475 cells migrating individually into the scratch wounds shown Fig. 5A. Boxplot shows the median, 25th, and 75th percentile with whiskers reaching the last data point; dots, fields of view; *n* = 47 (WT), 47 (KO), 50 (ΔIFBD), 24 (WT+PST) fields of view; *N* = 5 (WT, KO, ΔIFBD), 3 (WT+PST). Two-tailed *t*-test; †*P* < 0.001. **(C)** Quantification of N-cadherin (N-cad) and vimentin in indicated SNU-475 cell lines by immunoblotting. GAPDH, loading control. Boxplots show relative band intensities normalized to GAPDH. The box represents the median, 25th, and 75th percentile with whiskers reaching the last data point; dots, individual experiments; *N* = 8. Two-tailed *t*-test; **P* < 0.05, ***P* < 0.01. **(D)** Spider plots with migration trajectories of WT, KO, ΔIFBD, and PST-treated WT (WT+PST) SNU-475 cells tracked during 16 hours of random migration; dots, the final position of each single

tracked cell. **(E)** Quantification of persistence indices of WT, KO, ΔIFBD, and PST-treated WT (WT+PST) SNU-475 cells shown in (D). Boxplot shows the median, 25th, and 75th percentile with whiskers reaching the last data point; dots, individual cells; *n* = 15 (WT), 15 (KO), 19 (ΔIFBD), 14 (WT+PST); *N* = 3. **(F)** Quantification of average protrusion frequency of WT, KO, ΔIFBD, and PST-treated WT (WT+PST) SNU-475 cells during random and EGF-guided (directed) migrations. Boxplot shows the median, 25th, and 75th percentile with whiskers reaching the last data point; dots, individual cells; *n* (random) = 23 (WT), 21 (KO), 21 (ΔIFBD), 20 (WT+PST) cells; *n* (directed) = 15 (WT), 15 (KO), 17 (ΔIFBD), 14 (WT+PST) cells; *N* = 3. Two-tailed *t*-test; **P* < 0.05, ******$P$ < 0.01; **†**$P$ < 0.001. **(G)** Quantification of migration velocity of WT, KO, ΔIFBD, and PST-treated WT (WT+PST) SNU-475 cells during random and EGF-guided (directed) migrations. Boxplot shows the median, 25th, and 75th percentile with whiskers reaching the last data point; dots, individual cells; *n* (random) = 23 (WT), 21 (KO), 21 (ΔIFBD), 20 (WT+PST) cells; *n* (directed) = 15 (WT), 15 (KO), 17 (ΔIFBD), 14 (WT+PST) cells; *N* = 3. Two-tailed *t*-test; **†**$P$ < 0.001.

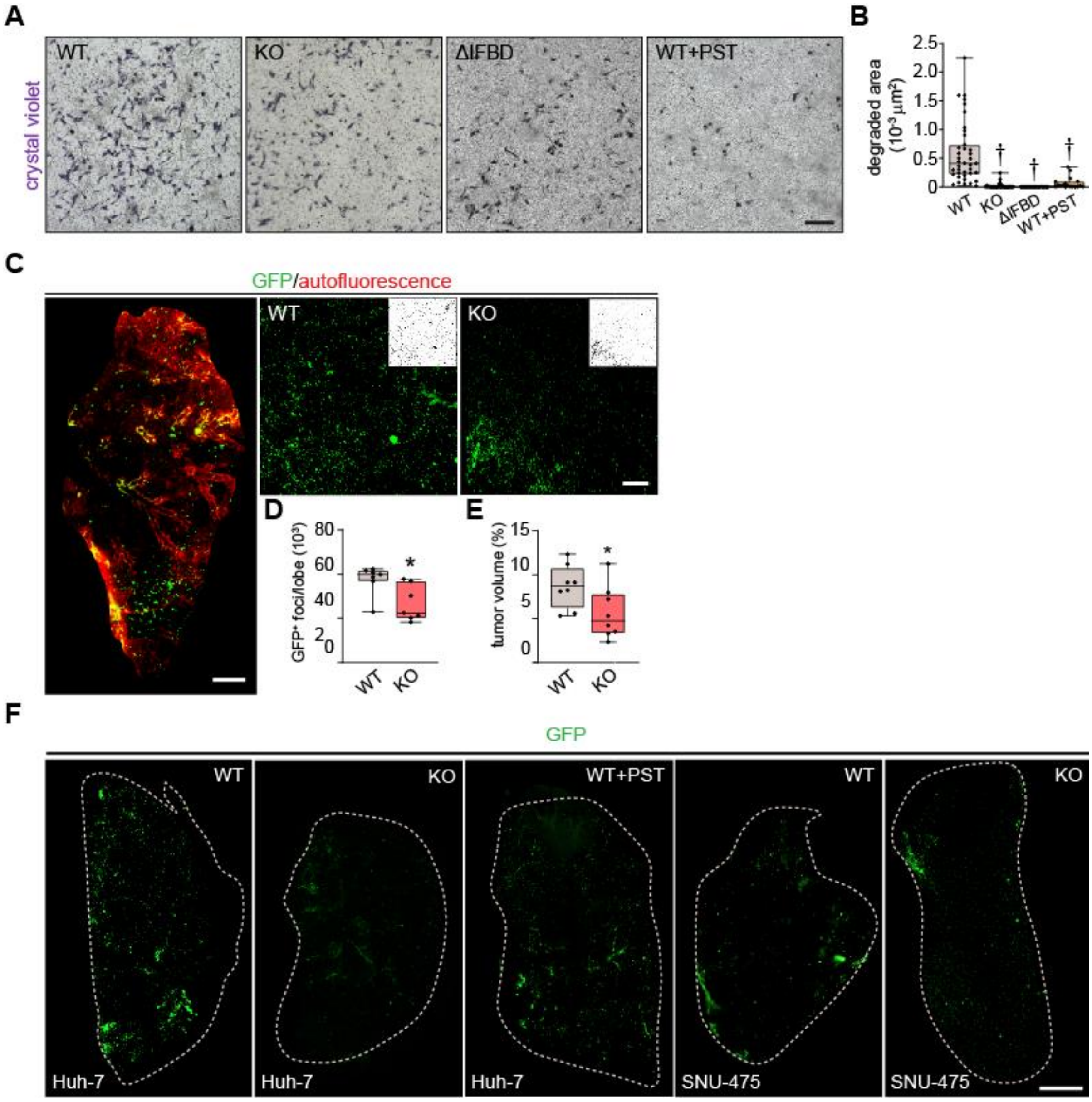


**Figure 6—figure supplement 1. Plectin inactivation reduces invasiveness of HCC cells. (A)** Representative images of crystal violet-stained WT, KO, ΔIFBD, and PST-treated WT (WT+PST) SNU-475 cells invading in Matrigel transwell assay shown in Fig. 6C. Scale bar, 300 µm. **(B)** Quantification of gelatin area degraded by WT, KO, ΔIFBD, and PST-treated WT (WT+PST) SNU-475 cells shown in Fig. 6D. Boxplot shows the median, 25th, and 75th percentile with whiskers reaching the last data point; dots, individual cells; *n* = 49 (WT), 36 (KO), 25 (ΔIFBD), 29 (WT+PST); *N* = 5 (WT), 4 (KO), 3 (IFBD, ΔWT+PST). Two-tailed *t*-test; †*P* < 0.001. **(C)** The 5-week-old NSG mice were injected (t.v.i.) with indicated RedFLuc-GFP-expressing SNU-475 cells. Representative lattice light sheet fluorescence image of CUBIC-cleared lung lobe immunolabeled with antibodies against GFP (green). Autofluorescence visualizing the lobe structures is shown in red. Scale bar, 1 mm. Representative magnified images from lung lobes with GFP-positive WT and KO nodules. Insets, segmented binary

masks of GFP-positive metastatic nodules. Scale bar, 200 µm. **(D,E)** Boxplots show metastatic load in the lungs expressed as the number (D) and relative volume (E) of GFP-positive ($GFP^+$) nodules indicated in (C). The box represents the median, 25th, and 75th percentile with whiskers reaching the last data point; dots, lung lobes; $n$ = 8 lung lobes; $N$ = 4. Two-tailed $t$-test; *$P < 0.05$ **(F)** Representative lattice light sheet fluorescence images of CUBIC-cleared lung lobes from (C) and Figure 6I, immunolabeled with antibodies against GFP (green). Dashed line, lobe contour. Scale bar, 1500 µm.

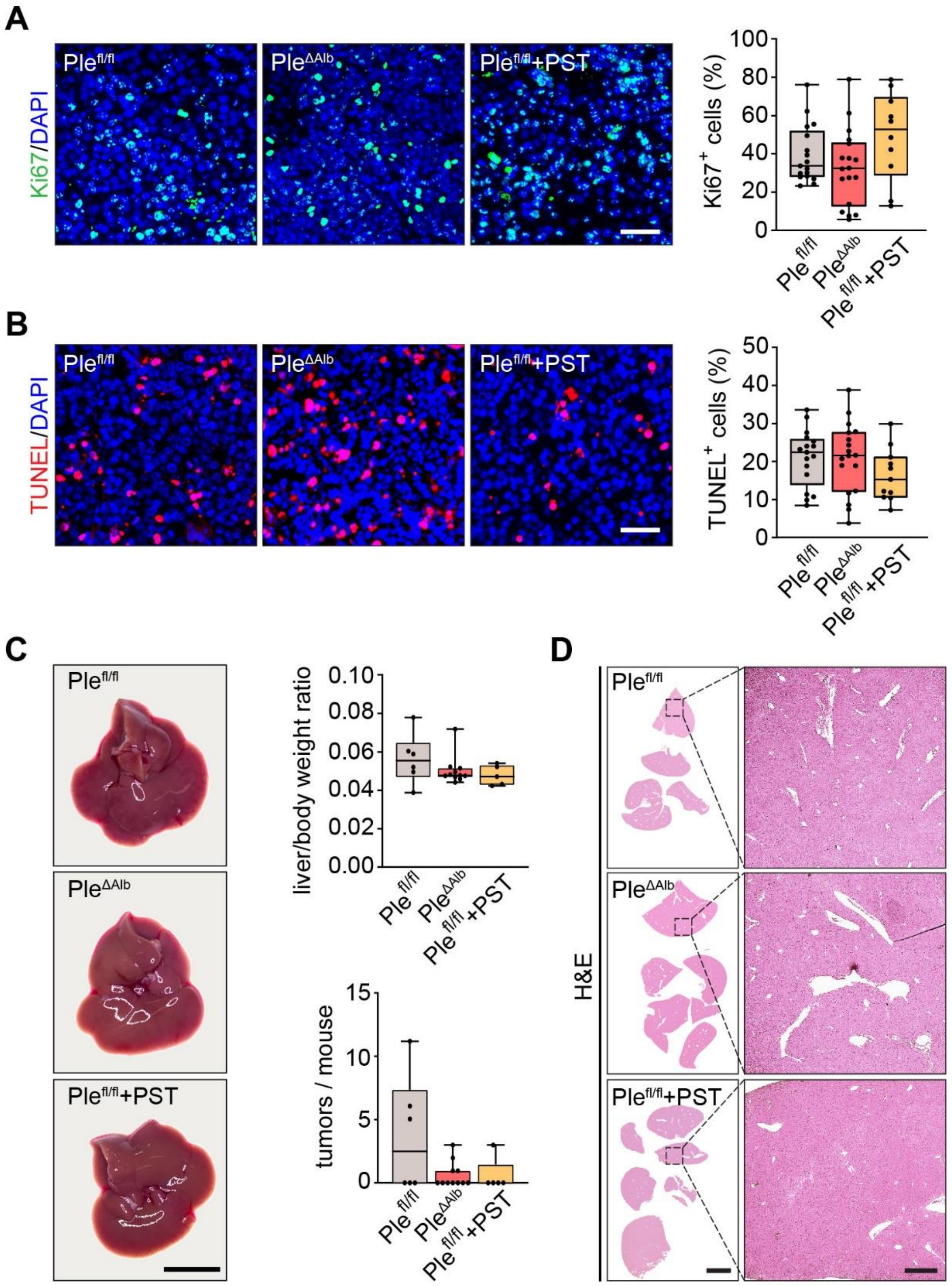


**Figure 7—figure supplement 1. Effect of plectin inactivation on HCC proliferation, apoptosis, and development. (A,B)** Representative images of Myc;sgTp53-induced liver tumor sections from *Ple*$^{fl/fl}$, *Ple*$^{\Delta Alb}$, and PST-treated *Ple*$^{fl/fl}$ (*Ple*$^{fl/fl}$ +PST) mice shown in Fig. 7B,C immunolabeled for Ki67 (green; A) and TUNEL (red;B). Nuclei, DAPI (blue). Scale bar, 50 µm. Quantification (percentage) of Ki67-positive cells (A) and TUNEL-positive cells (B). Boxplot shows the median, 25th and 75th percentile with whiskers reaching the last data point; dots, individual tumors; *n* = 17 (*Ple*$^{fl/fl}$), 18 (*Ple*$^{\Delta Alb}$), 10 (*Ple*$^{fl/fl}$+PST). **(C,D)** Myc;sgTp53 HCC was induced as before (see Fig. 7A,B) in 8-week-old *Ple*$^{fl/fl}$ and

*Ple*$^{\Delta Alb}$ female mice. *Ple*$^{fl/fl}$ mice were kept either untreated or every second day provided with orogastric gavage of plecstatin (*Ple*$^{fl/fl}$+PST). Animals were sacrificed 8 weeks post-induction. Representative images of *Ple*$^{fl/fl}$, *Ple*$^{\Delta Alb}$, and *Ple*$^{fl/fl}$+PST livers. Scale bar, 1 cm. Boxplots show tumor burden in the livers expressed as the liver/body weight ratio (upper graph) and number of tumors per mouse (lower graph). The box represents the median, 25th, and 75th percentile with whiskers reaching the last data point; dots, mice; *N* = 6 (*Ple*$^{fl/fl}$), 13 (*Ple*$^{\Delta Alb}$), 5 (*Ple*$^{fl/fl}$ + PST). **(D)** Representative images of H&E-stained *Ple*$^{fl/fl}$, *Ple*$^{\Delta Alb}$, and *Ple*$^{fl/fl}$+PST sections of livers shown in (C). Boxed areas, x12 images. Scale bars, 5 and 1 mm (boxed areas).

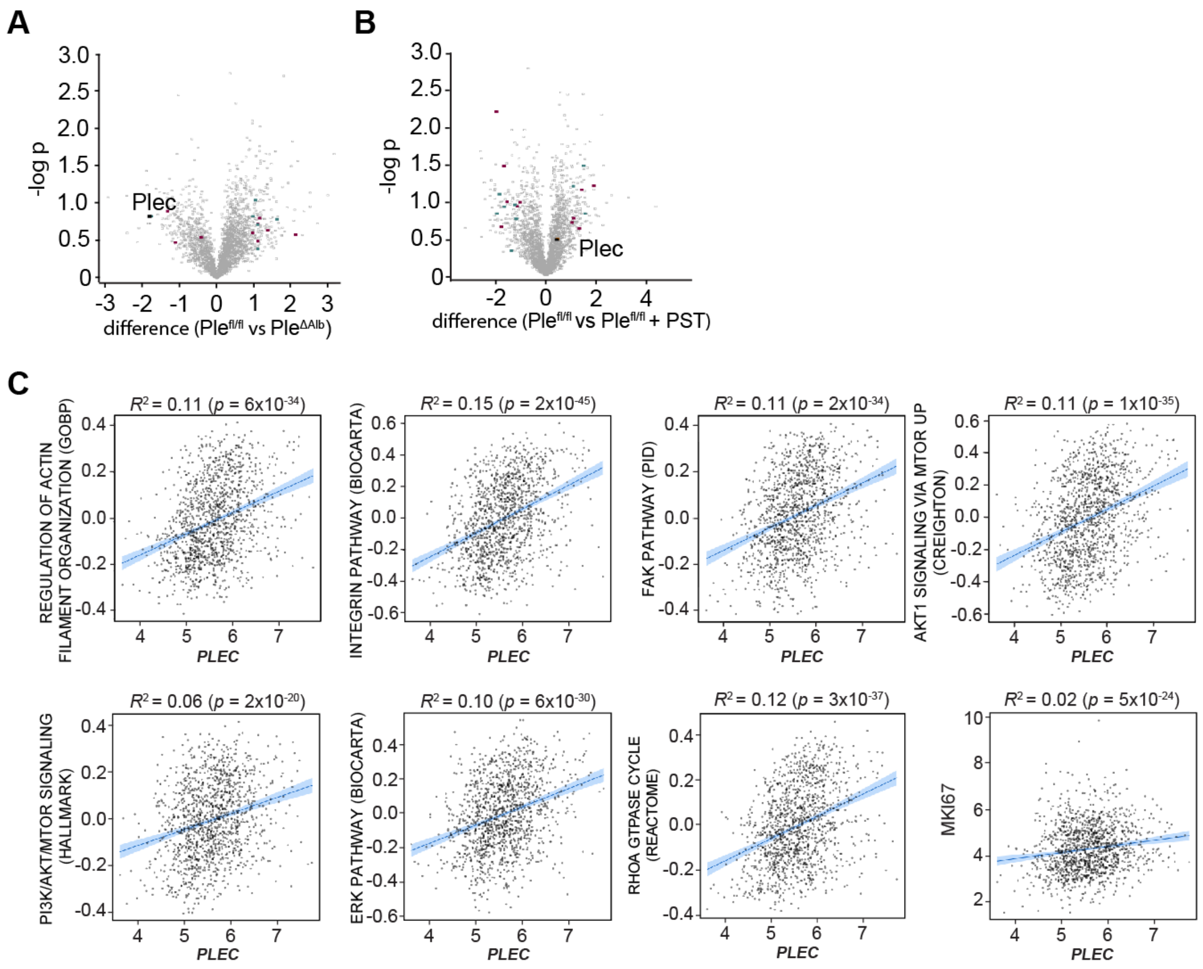


**Figure 7—figure supplement 2. Plectin-related expression signatures HCC from animal models and patients. (A,B)** Volcano plots show the fold change vs. *p*-value of differentially expressed proteins in livers of $Ple^{fl/fl}$ vs. $Ple^{\Delta Alb}$ (A) and $Ple^{fl/fl}$ vs. $Ple^{fl/fl}$ + PST (B) Myc;sgTp53-treated mice (see also Fig. 7B). Colored dots represent differentially expressed proteins identified in canonical signaling pathways shown in Fig. 7D. **(C)** Scatter plots show the correlation of *plectin* (*PLEC*) mRNA expression with indicated expression signatures. Line, linear regression line; blue area, 95% confidence intervals. Coefficient of determination ($R^2$) and $P$ values are indicated above the graphs.

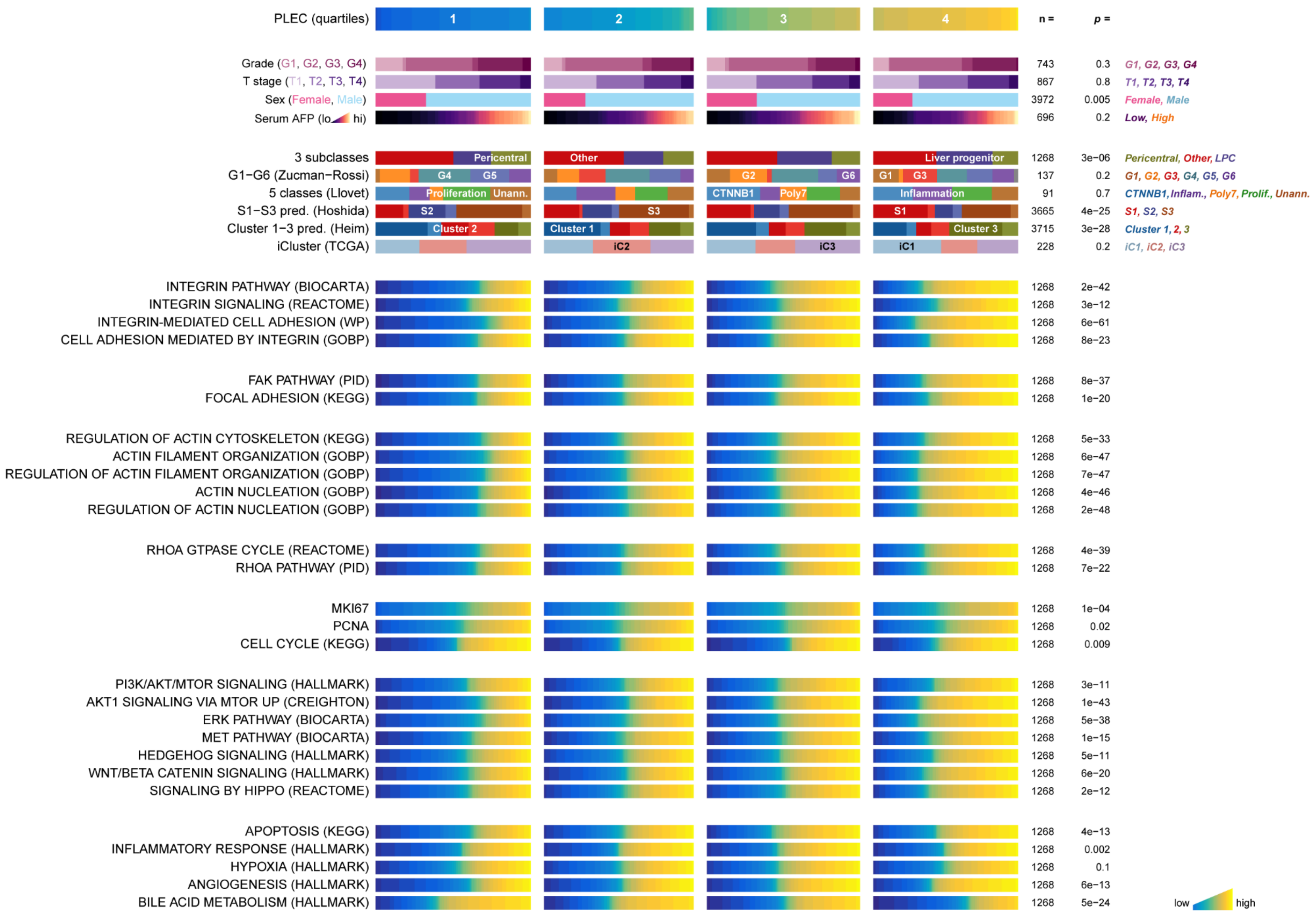
PLEC (quartiles)
1
2
3
4
n =
p =
Grade (G1, G2, G3, G4) 743 0.3 G1, G2, G3, G4
T stage (T1, T2, T3, T4) 867 0.8 T1, T2, T3, T4
Sex (Female, Male) 3972 0.005 Female, Male
Serum AFP (lo hi) 696 0.2 Low, High
3 subclasses 1268 3e−06 Pericentral, Other, LPC
Pericentral
Other
Liver progenitor
G1−G6 (Zucman−Rossi) 137 0.2 G1, G2, G3, G4, G5, G6
G4
G5
G2
G6
G1
G3
5 classes (Llovet) 91 0.7 CTNNB1, Inflam., Poly7, Prolif., Unann.
Proliferation
Unann.
CTNNB1
Poly7
Inflammation
S1−S3 pred. (Hoshida) 3665 4e−25 S1, S2, S3
S2
S3
S1
Cluster 1−3 pred. (Heim) 3715 3e−28 Cluster 1, 2, 3
Cluster 2
Cluster 1
Cluster 3
iCluster (TCGA) 228 0.2 iC1, iC2, iC3
iC2
iC3
iC1
INTEGRIN PATHWAY (BIOCARTA) 1268 2e−42
INTEGRIN SIGNALING (REACTOME) 1268 3e−12
INTEGRIN-MEDIATED CELL ADHESION (WP) 1268 6e−61
CELL ADHESION MEDIATED BY INTEGRIN (GOBP) 1268 8e−23
FAK PATHWAY (PID) 1268 8e−37
FOCAL ADHESION (KEGG) 1268 1e−20
REGULATION OF ACTIN CYTOSKELETON (KEGG) 1268 5e−33
ACTIN FILAMENT ORGANIZATION (GOBP) 1268 6e−47
REGULATION OF ACTIN FILAMENT ORGANIZATION (GOBP) 1268 7e−47
ACTIN NUCLEATION (GOBP) 1268 4e−46
REGULATION OF ACTIN NUCLEATION (GOBP) 1268 2e−48
RHOA GTPASE CYCLE (REACTOME) 1268 4e−39
RHOA PATHWAY (PID) 1268 7e−22
MKI67 1268 1e−04
PCNA 1268 0.02
CELL CYCLE (KEGG) 1268 0.009
PI3K/AKT/MTOR SIGNALING (HALLMARK) 1268 3e−11
AKT1 SIGNALING VIA MTOR UP (CREIGHTON) 1268 1e−43
ERK PATHWAY (BIOCARTA) 1268 5e−38
MET PATHWAY (BIOCARTA) 1268 1e−15
HEDGEHOG SIGNALING (HALLMARK) 1268 5e−11
WNT/BETA CATENIN SIGNALING (HALLMARK) 1268 6e−20
SIGNALING BY HIPPO (REACTOME) 1268 2e−12
APOPTOSIS (KEGG) 1268 4e−13
INFLAMMATORY RESPONSE (HALLMARK) 1268 0.002
HYPOXIA (HALLMARK) 1268 0.1
ANGIOGENESIS (HALLMARK) 1268 6e−13
BILE ACID METABOLISM (HALLMARK) 1268 5e−24
low
high

**Figure 7—figure supplement 3. Plectin signature in HCC patients.** The graph shows the association of *plectin* (*PLEC*) mRNA expression with indicated clinicopathological parameters, molecular classifications, and signature pathways among patients grouped into quartiles of PLEC expression. The analysis is based on data from gene set variation analysis (GSVA) used to produce quantitative enrichment scores for all gene sets from msigdb in pooled and batch-adjusted data. *P* values represent the result of a *chi*-square test (for categorical data) or analysis of variance (for numerical data such as gene signature expression levels).

**Supplemental Video Legends**

**Video 1.** Representative video of WT and KO SNU-475 cells invading the matrigel. Time-lapse covers total 21 h with frame taken every 15 min (∼15 min elapsed time per frame of the movie). Scale bar, 200 µm. The fixed and immunolabeled cells from the endpoint of this experiment are shown in Fig. 6E.

**Supplementary file 1**. Table of patients clinical data.

| **Sample ID** | **Etiology**<br>0 = unknown<br>1 = ALD/ASH<br>2 = NAFLD/NASH<br>3 = HBV<br>4 = HCV<br>5 = Other | **Age** [years] | **Sex**<br>0 = male<br>1 = female |
|---|---|---|---|
| MAP_1_HCC | 4 | 78.5 | 1 |
| MAP_10_HCC | 4 | 51.4 | 0 |
| MAP_11_HCC | 2 | 79.0 | 0 |
| MAP_12_HCC | 1 | 86.2 | 0 |
| MAP_13_HCC | 1 | 69.2 | 0 |
| MAP_14_HCC | 3 + 4 | 49.5 | 0 |
| MAP_15_HCC | 1 | 83.9 | 0 |
| MAP_16_HCC | 4 | 59.2 | 1 |
| MAP_17_HCC | 4 | 54.8 | 0 |
| MAP_18_HCC | 1 + 4 | 68.4 | 0 |
| MAP_19_HCC | 1 + 4 | 56.5 | 1 |
| MAP_2_HCC | 2 + 5 | 61.8 | 1 |
| MAP_20_HCC | 1 + 4 | 48.2 | 0 |
| MAP_21_HCC | 2 + 5 | 63.5 | 1 |
| MAP_3_HCC | 1 | 67.5 | 0 |
| MAP_4_HCC | 1 | 71.9 | 0 |
| MAP_5_HCC | 0 | 66.1 | 0 |
| MAP_6_HCC | 1 | 83.0 | 0 |
| MAP_7_HCC | 0 | 63.3 | 1 |
| MAP_8_HCC | 2 | 68.6 | 0 |
| MAP_9_HCC | 3 | 57.9 | 0 |

**Supplementary file 2.** List of antibodies used in this study.

| **Primary antibodies** | | | | | |
|---|---|---|---|---|---|
| **Antigen** | **Clone** | **Manufacturer** | **Cat. number** | **Application** | **Dilution** |
| rabbit anti-Akt | polyclonal | Cell Signaling | 9272 | WB | 1:1000 |
| rabbit anti-pAkt (Ser473) | D9E | Cell Signaling | 4060 | WB | 1:2000 |
| mouse anti-E-cadherin | 36/E | BD Bioscience | 610181 | WB | 1:4000 |
| mouse anti-Erk2 | D-2 | Santa Cruz | sc-1674 | WB | 1:200 |
| rabbit anti-pErk1/2 (Thr202/Tyr204) | D13.14.4E | Cell Signaling | 4370 | WB | 1:1000 |
| mouse anti-FAK | 77/FAK Ruo | BD Bioscience | 610088 | WB | 1:500 |
| rabbit anti p-FAK (Tyr397) | polyclonal | Thermo-Fisher | 44-624G | WB | 1:1000 |
| rabbit anti-GAPDH | FF26A | Sigma | G9545 | WB | 1:20.000 |
| rabbit anti-GFP | polyclonal | Invitrogen | A-11122 | immunofluorescence | 1:100 |
| rabbit anti-ILK | 4G9 | Cell Signaling | 3856 | WB | 1:1000 |
| rabbit anti- αV integrin | EPR16800 | Abcam | ab179475 | WB | 1:5000 |
| mouse anti-β1 integrin | A-4 | Santa Cruz | sc-374429 | WB | 1:500 |
| rabbit anti-Ki67 | SP6 | GeneTex | GTX16667 | immunofluorescence | 1:100 |
| rabbit anti-N-cadherin | polyclonal | Abcam | ab18203 | WB | 1:1000 |
| mouse anti-paxillin | 349 | BD Bioscience | 612405 | WB | 1:1000 |
| rabbit anti-PI3K p85 | 19H8 | Cell Signaling | 4257 | WB | 1:1000 |
| rabbit anti-pPI3K p85/p55 (Tyr458/Tyr199) | polyclonal | Cell Signaling | 4228 | WB | 1:1000 |
| guinea pig anti-plectin | polyclonal | Progen | GP21 | WB | 1:1000 |
| | | | | immunofluorescence | 1:250 |
| mouse anti-talin | 8d4 | Sigma | T3287 | WB | 1:1000 |
| mouse anti-vimentin | RV202 | Santa Cruz | sc-32322 | WB | 1:500 |
| rabbit anti-vimentin | polyclonal | GeneTex | GTX100619 | immunofluorescence | 1:200 |
| mouse anti-vinculin | VIN-11-5 | Sigma | V4505 | WB | 1:500 |
| | | | | immunofluorescence | 1:250 |
| **Secondary antibodies** | | | | | |
| **Name** | **Manufacturer** | | **Cat. number** | **Application** | **Dilution** |
| anti-mouse AF-488 | Jackson ImmunoResearch | | 715-545-150 | immunofluorescence | 1:500 |
| anti-rabbit AF-488 | Jackson ImmunoResearch | | 711-545-152 | immunofluorescence | 1:500 |
| anti-rabbit AF-594 | Life Tech | | A11037 | immunofluorescence | 1:1000 |
| anti-rabbit AF-647 | Jackson ImmunoResearch | | | immunofluorescence | 1:500 |
| anti-guinea pig AF-488 | Jackson ImmunoResearch | | 706-545-148 | immunofluorescence | 1:500 |
| anti-mouse IgG IRDye® 680RD | Licor | | 926-68072 | WB | 1:20,000 |
| anti-mouse IRDye® 800CW | Licor | | 926-32212 | WB | 1:20,000 |
| anti-rabbit IRDye® 680RD | Licor | | 926-68073 | WB | 1:20,000 |
| anti-rabbit IgG IRDye® 800CW | Licor | | 926-32213 | WB | 1:20,000 |
| anti-guinea pig IRDye® 680RD | Licor | | 926-68077 | WB | 1:20,000 |
| anti-mouse-IgG-HRP | Jackson ImmunoResearch | | 115-035-146 | WB | 1:5000 |
| anti-rabbit-IgG-HRP | Jackson ImmunoResearch | | 111-035-044 | WB | 1:5000 |
| anti-GuineaPig-IgG-HRP | Sigma | | A7289 | WB | 1:5000 |
